# Spinal motoneurons respond aberrantly to serotonin in a rabbit model of cerebral palsy

**DOI:** 10.1101/2023.04.05.535691

**Authors:** E. J. Reedich, L.T. Genry, P.R. Steele, E. Mena Avila, L. Dowaliby, A. Drobyshevsky, M. Manuel, K. A. Quinlan

## Abstract

Cerebral palsy (CP) is caused by a variety of factors that damage the developing central nervous system. Impaired motor control, including muscle stiffness and spasticity, is the hallmark of spastic CP. Rabbits that experience hypoxic-ischemic (HI) injury *in utero* (at 70-80% gestation) are born with muscle stiffness, hyperreflexia, and, as recently discovered, increased serotonin (5-HT) in the spinal cord. To determine whether serotonergic modulation of spinal motoneurons (MNs) contributes to motor deficits, we performed *ex vivo* whole cell patch clamp in neonatal rabbit spinal cord slices at postnatal day (P) 0-5. HI MNs responded to application of α-methyl 5-HT (a 5-HT_1_/5-HT_2_ receptor agonist) and citalopram (a selective 5-HT reuptake inhibitor) with hyperpolarization of persistent inward currents and threshold voltage for action potentials, reduced maximum firing rate, and an altered pattern of spike frequency adaptation while control MNs did not exhibit any of these responses. To further explore the differential sensitivity of MNs to 5-HT, we performed immunohistochemistry for inhibitory 5-HT_1A_ receptors in lumbar spinal MNs at P5. Fewer HI MNs expressed the 5-HT_1A_ receptor compared to age-matched controls. This suggests many HI MNs lack a normal mechanism of central fatigue mediated by 5-HT_1A_ receptors. Other 5-HT receptors (including 5-HT_2_) are likely responsible for the robust increase in HI MN excitability. In summary, by directly exciting MNs, the increased concentration of spinal 5-HT in HI rabbits can cause MN hyperexcitability, muscle stiffness, and spasticity characteristic of CP. Therapeutic strategies that target serotonergic neuromodulation may be beneficial to individuals with CP.

**Key points:**

- After prenatal hypoxia-ischemia (HI), neonatal rabbits that show hypertonia are known to have higher levels of spinal serotonin
- We tested responsivity of spinal motoneurons (MNs) in neonatal control and HI rabbits to serotonin using whole cell patch clamp
- MNs from HI rabbits showed a more robust excitatory response to serotonin than control MNs, including hyperpolarization of the persistent inward current and threshold for action potentials, larger post-inhibitory rebound, and less spike frequency adaptation
- Based on immunohistochemistry of lumbar MNs, fewer HI MNs express inhibitory 5HT_1A_ receptors than control MNs, which could account for the more robust excitatory response of HI MNs.
- These results suggest that after HI injury, the increased serotonin could trigger a cascade of events leading to muscle stiffness and altered motor unit development

## Introduction

Clinical features of spastic cerebral palsy (CP) include nonprogressive muscle/joint stiffness and hyperreflexia (Volpe et al., 2017). CP is caused by a variety of insults to the developing nervous system, often involving hypoxia, that cause lifelong motor deficits. Perinatal hypoxia-ischemia (HI) can occur with maternal infection, inflammation, placental insufficiency, and a difficult birth, and causes white matter injury, thinning of the corticospinal tract, cortical damage, and motor deficits, which are the hallmarks of CP (Derrick et al., 2004; Buser et al., 2010; Drobyshevsky and Quinlan, 2017). The pathophysiological mechanisms of these motor deficits are not well understood, so there is a paucity of evidence-based treatments for this condition. Many therapies for CP do not consistently result in improvements even though some approaches can be painful (Novak et al., 2013; Parkinson et al., 2013; Wimalasundera and Stevenson, 2016). Other common treatment strategies, like botulinum neurotoxin injection, diminish symptoms without correcting chronic underlying neurological issues. A better understanding of the neurological causes of spasticity and dystonia in CP would make it possible to develop evidence-based therapies and/or pharmaceutical treatments to prevent or ameliorate this debilitating symptom.

An animal model of CP that shows clear motor phenotypes is needed to make advancements. New Zealand White rabbits (perinatal brain developers) show prominent motor deficits consistent with spastic CP after prenatal HI (at 70-80% gestation). These deficits include muscle stiffness (Derrick et al., 2004), loss of neurons in cortical layers 3 and 5, white matter injury, thinning of the corticospinal tract (Buser et al., 2010), cell death in the spinal cord and decreased numbers of spinal MNs (Drobyshevsky and Quinlan, 2017), increased sarcomere length, decreased muscle mass and hyperreflexia (Synowiec et al., 2019), and higher firing frequencies and more sustained firing in affected spinal MNs (Steele et al., 2020). In stark contrast, rodents (postnatal brain developers) show only mild motor deficits, even after severe brain injuries that are used to mimic CP-causing events (Rice et al., 1981; Fan et al., 2005; Cavarsan et al., 2019). This discrepancy is likely due to a mismatch in neural development, since humans (like rabbits) are perinatal brain developers (Dobbing and Sands, 1973, 1979; Clancy et al., 2007; Semple et al., 2013). Thus, to better mimic outcomes of perinatal brain injuries in humans, rodents are usually injured at one week of age (Rice et al., 1981). However, rodents are already learning to walk at this developmental stage (Fox, 1965). Rabbit neuromuscular development is better matched to humans, making them a more effective model for CP research.

Serotonin (5-HT) is generally thought of as a neurotransmitter and neuromodulator associated with anxiety and depression, but it is also critically important during development. Developmental disruption in 5-HT signaling is associated with neurological disorders including autism, Rett syndrome, Down’s syndrome and, more recently, CP (Bar-Peled et al., 1991; Whittle et al., 2007; Bellot et al., 2014; Yang et al., 2014; De Filippis et al., 2015; Drobyshevsky et al., 2015; Muller et al., 2016; Wirth et al., 2017). Typically, descending serotonergic tracts from the rostroventral medulla’s raphe nucleus are among the first to arrive in the spinal cord (at 9 weeks post conception in humans (Sundström et al., 1993)), and during periods of activity, the spinal cord is bathed in 5-HT through volume transmission (Bunin and Wightman, 1999; Hentall et al., 2006). The developing fetal spinal cord is exposed to 5-HT through multiple sources: the fetal brainstem raphe nucleus, the maternal raphe nucleus, and the placenta (Bonnin and Levitt, 2011). In humans, the concentration of 5-HT in the spinal cord progressively increases until reaching a developmental peak at 1-3 years of age (Lewinsohn et al., 1980; Lacković et al., 1988). Spinal 5-HT is elevated threefold in hypertonic HI rabbits at birth (Drobyshevsky et al., 2015). The impact of increased spinal 5-HT during development is not known. Since 5-HT affects development of MN excitability (Bou-Flores et al., 2000; Fricker et al., 2005; Mogha et al., 2012), we investigated the link between serotonergic neurotransmission and MN excitability in the HI rabbit model of spastic CP.

We tested the hypothesis that serotonergic neuromodulation causes MN hyperexcitability in rabbits subjected to HI *in utero* (“HI rabbits”) with the goal of identifying more effective targets for treatment of spastic CP. Muscle stiffness and hyperreflexia consistent with spastic CP are prominent features of many (but not all) rabbit kits after HI injury (Shi et al., 2021). Typically, 50% of HI rabbit kits have motor deficits (referred to as “HI affected”), while the other half of the litter does not have any motor deficits despite having experienced HI (referred to as “HI unaffected”). Spinal 5-HT concentration is significantly higher in HI affected kits compared to both control and HI unaffected kits (Drobyshevsky et al., 2015). Increased spinal 5-HT is sufficient to cause hypertonia, since intrathecal application of 5-HT increased joint stiffness in HI unaffected kits, but not in HI affected kits, which already had high levels of 5-HT and joint stiffness; conversely, inhibition of 5-HT_2_/activation of 5-HT_1_ receptors with methysergide decreased stiffness in HI-affected kits but had no effect in HI unaffected kits (Drobyshevsky et al., 2015). Intrathecal application of these drugs could have multiple effects in the spinal cord, where many neurons are responsive to 5-HT. In this study, we directly tested MN responsiveness to 5-HT with the 5-HT_1/2_ receptor agonist α-methyl 5-HT in the presence of the reuptake inhibitor citalopram, which would prolong the activity of endogenous 5-HT in the spinal cord (Maclean et al., 1998; Pearlstein et al., 2005). To further understand the receptors mediating these effects, we investigated 5-HT_1A_ receptor expression in spinal MNs via immunohistochemistry. We demonstrate here that HI MNs-from both HI affected and HI unaffected kits-respond to 5-HT receptor activation with a robust increase in excitability while control MNs have a very modest response. This may be due to reduced expression of inhibitory 5-HT_1A_ receptors and/or enhanced expression of 5-HT_2_ type receptors in HI MN pools. We conclude that elevated spinal 5-HT could contribute to development of spasticity by direct activation of MNs after perinatal injury to the central nervous system.

## Methods

### Hypoxia-ischemia (HI) surgery

All rabbits were used according to the University of Rhode Island’s Animal Care and Use Committee guidelines. Pregnant New Zealand White rabbits (bred in house or ordered from Charles River Laboratories, Inc, Wilmington MA), underwent HI procedures as described in (Derrick et al., 2004). Briefly, at ~70-80% gestation (day 22-26 of 31.5 days typical gestation) dams were premedicated with ketamine and xylazine (25 mg/kg and 3.75 mg/kg, respectively), anesthetized with isoflurane (1-3%; inhaled via V gel tube), and if necessary, treated with buprenorphine (0.01 - 0.24 mg/kg). Indications for treatment with buprenorphine include elevated heart rate (over 220 BPM); prolonged rapid, shallow breathing/panting; elevated CO_2_ levels (due to panting). The left or right femoral artery was isolated, incised, and a Fogarty balloon catheter was inserted and advanced to the level of the bifurcation of the descending aorta. The balloon catheter was inflated for 40 minutes, occluding blood flow to the uterine arteries and causing HI in the fetuses. After the procedure, dams recovered and later gave birth to kits with HI injuries. Categorization of whether the kits were affected or unaffected by HI was based on a modified Ashworth scale (as described in Derrick et al., 2004). Sham kits were born to dams that underwent the same procedures but without insertion and/or inflation of the catheter. Control kits were rabbits born without any prenatal interventions. Since there were no observable differences between sham and control groups in any of the properties measured in this study, they were grouped together here. Thus, the rabbits referred to as “control” in the remainder of this manuscript are a mix of surgical sham and unoperated naive control kits.

### Patch clamp electrophysiology

Whole cell patch clamp was performed similarly to our previously published work (Quinlan et al., 2011; Steele et al., 2020). Briefly, transverse spinal cord slices were obtained at 350 μm thickness using a Leica VT1200S vibratome. Immediately after slicing, sections were incubated for one hour at 30 °C in oxygenated (95% O_2_ and 5% CO_2_) artificial cerebrospinal fluid (aCSF). Recording was performed with aCSF containing 111 mM NaCl, 3.09 mM KCl, 25.0 mM NaHCO_3_, 1.10 mM KH_2_PO_4_, 1.26 mM MgSO_4_, 2.52 mM CaCl_2_, and 11.1 mM glucose at a perfusion rate of 2.5 mL/min. Whole cell patch electrodes (1-3 MΩ) contained 138 mM K-gluconate, 10 mM HEPES, 5 mM ATP-Mg, 0.3 mM GTP-Li and 150 μM 3000 MW Texas Red dextran. Persistent inward currents (PICs) were measured in voltage clamp mode with holding potential of −90 mV and depolarizing voltage ramps of both 22.5 mV/s and 11.25 mV/s bringing the cell to 0 mV in 4 s, and 8 s respectively, and then back to the holding potential in the following 4 or 8 seconds. In current clamp mode, action potential (AP) characteristics including threshold voltage (when rate of rise exceeded 10 V/s), overshoot past 0 mV, AP duration (at half amplitude between threshold and overshoot), and rate of rise and rate of fall (peak and trough of the first derivative of the voltage trace), as well as frequency-current relationships were obtained from current ramps. Hyperpolarizing and depolarizing current steps (1 s each) were used to test the hyperpolarization-activated current (I_h_; sag/rebound), threshold voltage, and spike frequency adaptation (SFA). To measure sag and rebound, the voltage was measured at the start and end of a hyperpolarizing current step, and after the cessation of the hyperpolarizing step. Sag voltage was the initial trough minus the final plateau voltage from the step, while rebound voltage was the amplitude of depolarization after the cessation of the step compared to the baseline voltage before the step. If an action potential fired after the step, the threshold voltage was used as the ending voltage. When measuring sag and rebound, hyperpolarizing steps of −300 pA to −2 nA were used, depending on the input resistance of the neuron. To normalize the sag/rebound elicited from different amplitude pulses, we calculated it as a percent of voltage change without the contribution from sag which was simply the voltage change due to sag or rebound compared to the hyperpolarization of the voltage without sag (the difference between resting membrane potential before the step to the membrane potential at the end of the step, after sag had diminished). To measure SFA, we calculated the spike accommodation index (SAI) for both the primary (1°P), or initial phase of firing as defined by previous studies (Shinomoto et al., 2003) and the secondary phase (2°P) which comprises the next few hundreds of milliseconds. Other studies, which have used longer current pulses, have referred to this phase as the early phase, which is followed by a late phase for the following tens of seconds to minutes (Granit et al., 1963; Kernell, 1965; Kernell and Monster, 1982; Smith and Brownstone, 2022). Here we focused on only primary and secondary phases, using 1 s current pulses. The SAI for the primary range was calculated using the first interspike interval (ISI) compared to the fourth ISI. The secondary range began with the fifth ISI divided by the last interspike interval from the same step (1°SAI=ISI_First_/ISI_4th_; 2°SAI=ISI_5th_/ISI_Last_). SAIs less than 1 indicate deceleration of the firing rate in response to a constant current (classical SFA), while SAIs greater than 1 indicate acceleration in the firing rate in the same conditions. Regarding neuron selection, neurons were targeted in the motoneuron (MN) pools mainly from cervical and lumbar regions of the spinal cord and were removed from the data set if their resting membrane potential was more depolarized than −35 mV at break in, or the neuron depolarized more than 10 mV after 5-HT application.

### Drug application

For each MN, once all properties were recorded in standard aCSF (see above), serotonergic drugs were bath applied and all electrophysiological parameters were recorded again. Either 10 μM serotonin (5-HT), or 0.3 μM α-methyl 5-HT in combination with 10 μM citalopram was applied. These drugs are hereafter referred to as the 5-HT cocktail. Citalopram was included because of the previous finding that serotonin transporter (*SERT*) expression is upregulated in HI affected rabbit spinal cords (Drobyshevsky et al., 2015). Since the perfusion rate was 2.5 mL/min and the tubing plus bath volume equaled 10 mL, all recordings were made 20 minutes after drug application, when the volume of the bath and tubing had been replaced roughly five times over.

### Immunostaining for 5-HT_1A_ receptors

#### Tissue harvests

P5 kits were deeply anesthetized by intraperitoneal injection of EUTHASOL^®^ Euthanasia Solution (pentobarbital sodium and phenytoin sodium; Patterson Veterinary, cat. # 07-805-9296). Transcardial perfusion was performed using chilled 0.01 M phosphate buffered saline (PBS; pH 7.4) until tissue blanching was observed, then alkaline paraformaldehyde (PFA; 4% in PBS) was infused at a constant flow rate of approximately 5-8 mL/min (depending on rabbit kit size). Perfused spinal cords were isolated and post-fixed in alkaline PFA (4% in PBS) for ~24-72 hrs. Fixed spinal cords were then washed with PBS and cryoprotected in 30% sucrose (in PBS) until sinking was observed. Cryoprotected spinal cords were embedded in optimal cutting temperature (OCT) compound and frozen at −80 °C until cryostat sectioning. Transverse lower lumbar spinal cord cryosections were collected in serial at 30-μm thickness using a Leica CM1950; to avoid double counting motoneurons, every 15^th^ section was used for analysis.

#### Immunostaining protocol

Slides were thawed and then dried in an oven at 37 °C to improve section adherence. A hydrophobic barrier was drawn along the perimeter of each slide with an ImmEdge™ Pen (Vector Laboratories, cat. # H-4000) to improve reagent retention during immunostaining. Antigen retrieval was performed by submerging slides in hot antigen retrieval buffer (10 mM citric acid, 0.05% Tween 20, pH 6.0; 70-80 °C) for 10 minutes. After rinsing with PBS, sections were blocked in blocking buffer (5% donkey serum, 0.5% Triton X-100, and 0.5% glycine in PBS) for 1 hour at room temperature. Following blocking, sections were incubated in primary antibody solution containing goat-anti ChAT antibody (Millipore Sigma, cat. #AB144P; 1:100 in blocking buffer) overnight at 4 °C. Sections were washed in PBST (0.1% Tween-20 in PBS) and then incubated in secondary antibody solution consisting of donkey anti-goat-488 (IgG H+L) (Jackson ImmunoResearch, cat. # 705-545-147; 1:500 in blocking buffer) for 3 h at room temperature. After rinsing with PBST, sections were re-blocked in blocking buffer for 1 hour at room temperature then incubated in primary antibody solution containing mouse anti-5-HT_1A_ receptor (Millipore Sigma, cat. # MAB11041; 1:200 in blocking buffer) overnight at 4 °C. Slices were washed in PBST and then incubated in secondary antibody solution consisting of donkey anti-mouse-594 (IgG H+L) (Jackson ImmunoResearch, cat. # 715-585-150; 1:500 in blocking buffer) for 3 h at room temperature. Nuclei were labeled with 4’,6-Diamidino-2-Phenylindole (DAPI) and slides were mounted using Fluoromount mounting medium (Sigma, cat. # F4680).

#### Confocal imaging and analysis

Z-stack photomicrographs (z step: 1 μm) of lower lumbar (L4-L7) spinal cord ventral horns (VHs) were acquired at 10x magnification using a Nikon Eclipse Ti2 inverted confocal microscope. Maximum intensity projections of the green, red, and blue channels were generated for each z-stack and merged using ImageJ. For each resultant VH image, blinded quantification of ChAT-immunoreactive MNs (with discernable nuclei) and the subset of MNs expressing 5-HT_1A_ receptors was performed. The relative abundance (percent) of 5-HT_1A_ receptor-positive MNs for each rabbit was calculated by pooling 5-HT1A-positive MNs and total MNs from all analyzed VHs (total 5-HT_1A_-positive MNs ÷ total MNs × 100).

### Statistics

Statistical analyses were performed with GraphPad Prism 9.5.0 and Microsoft Excel. Two-tailed paired t-tests were performed in Excel for all cells that were tested before and after application of the 5-HT cocktail. All other tests were performed using GraphPad Prism. Normality was assessed either visually using QQ plots of residuals or using a Shapiro-Wilk normality test. Data sets with only two groups that were normally distributed and had equal variances were analyzed using an unpaired t-test to assess group differences. A two-way repeated measures ANOVA was used to account for the correlation between repeated measurements and to determine if the cells in the control and HI groups were responding differently to 5-HT application. No post-hoc tests were performed due to the primary interest in the interaction effect.

## Results

### Motoneuron persistent inward currents

Motoneuron (MN) electrophysiological properties were measured before and after application of the 5-HT cocktail. The response of MNs from HI kits to the 5-HT cocktail was different from that of control kits. Persistent inward currents (PICs) are long-lasting voltage-sensitive depolarizing sodium and calcium currents that enable self-sustained firing in MNs (Heckman et al., 2008, 2009). Onset voltage and amplitude of PICs from control MNs showed no significant changes in the presence of the 5-HT cocktail while HI MNs showed significant changes consistent with increased excitability (**Figure 1**). We present HI MNs as a single group, for simplicity, because both HI unaffected and HI affected MNs responded the same way: both groups showed hyperpolarization of PIC onset in the presence of 5-HT cocktail (HI unaffected MNs before v. after 5-HT, *P*=0.006; HI affected MNs before v. after 5-HT, *P*=0.010). The voltage at PIC peak, another measure of the voltage dependence of the PIC, was affected similarly: control MNs did not show any change with the 5-HT cocktail (before v. after 5-HT, *P*=0.515) while voltage at PIC peak became significantly hyperpolarized in HI MNs (before v. after 5-HT, *P*=0.003; HI affected only, *P*=0.043; HI unaffected only, *P*=0.044). In addition to voltage dependence of the PIC, the presence of 5-HT cocktail also changed PIC amplitude in HI but not control MNs (Figure 1D). In this case, the effect appears to be more modest (see Figure 1D; HI MNs before v. after 5-HT, *P*=0.023), and when HI affected and unaffected MNs were separated, neither group showed statistical significance with paired t-tests (HI affected only, *P*=0.066; HI unaffected only, *P*=0.234). Overall, hyperpolarization of the PIC, which increases MN excitability, was a robust feature of HI MN responses to the 5-HT cocktail and was absent in the control group. In fact, two-way repeated measures ANOVA revealed a significant interaction effect between surgical group (control or HI) and drug treatment (presence or absence of 5-HT cocktail) for PIC amplitude (F[1,19] = 4.55, *P*=0.046) and voltage at PIC peak (F[1,19] = 6.44, *P*=0.020), demonstrating that PICs of control and HI MNs are differently modulated by 5-HT. See **Table 1** for a complete list of parameters measured in this study.

**Figure 1:**
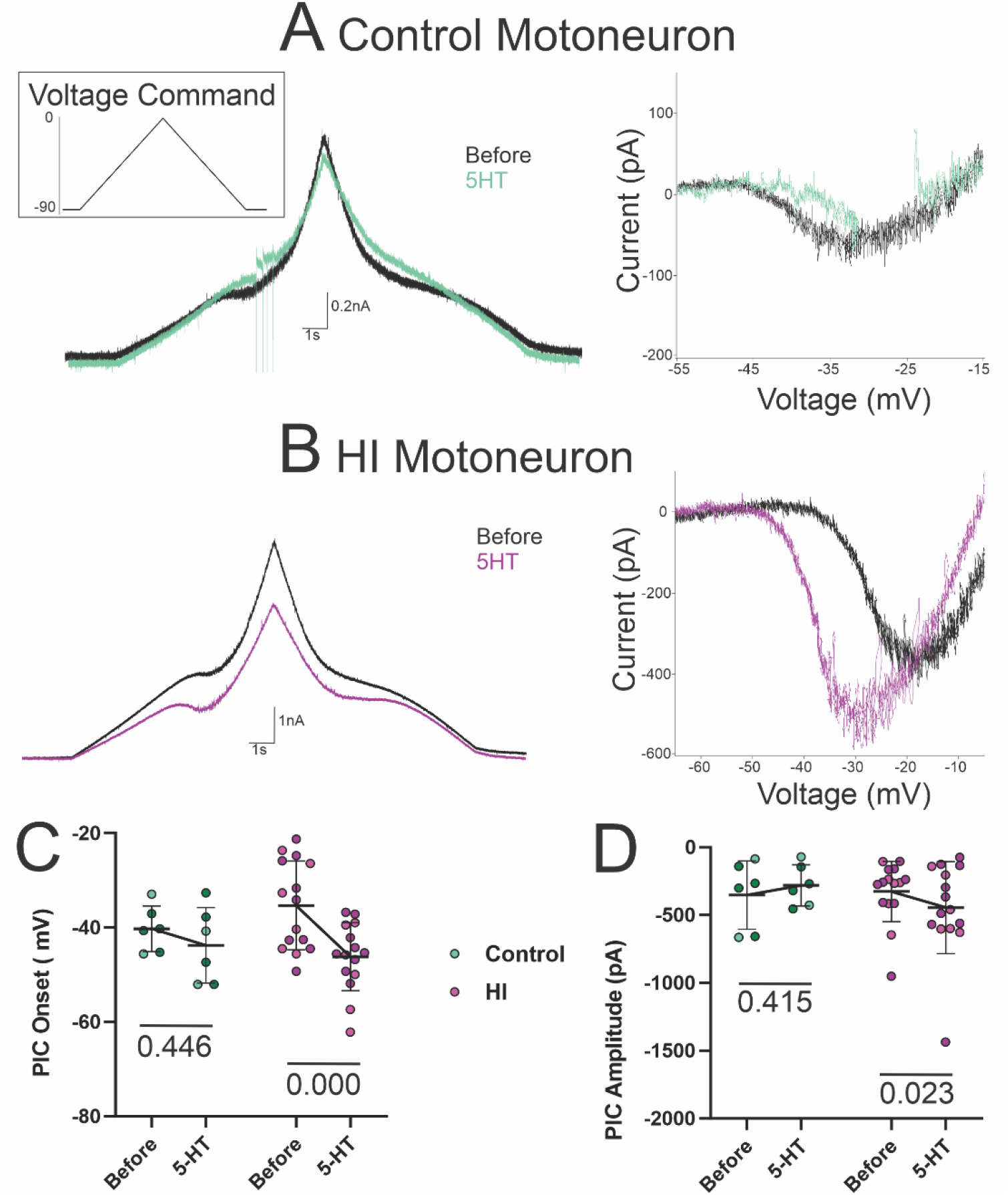
Persistent inward currents (PICs) were differently modulated by 5-HT cocktail in control and HI MNs. Examples of PICs recorded from control **(A)**, and HI **(B)** MNs evoked with voltage ramps (inset). PIC onset voltage (**C**) and PIC amplitude (**D**) were not significantly altered by 5-HT in control MNs, but both parameters were significantly altered by 5-HT in HI MNs. Data are represented as mean ± SD; N=6 control MNs and N=15 HI MNs (all before and after application of 5-HT cocktail). *P* values, displayed above pairs, are based on paired t-tests (before and after application of 5-HT cocktail). Kits constituting the sham subgroup (N = 4) of the control group are shaded dark green and kits constituting the motor-affected subgroup (N = 9) of the HI group are shaded dark pink. Lines connecting means before and after 5-HT visually represent the interaction effect of the two-way repeated measures ANOVA. This is seen by comparing the slopes of the lines between groups.

**Table 1:**
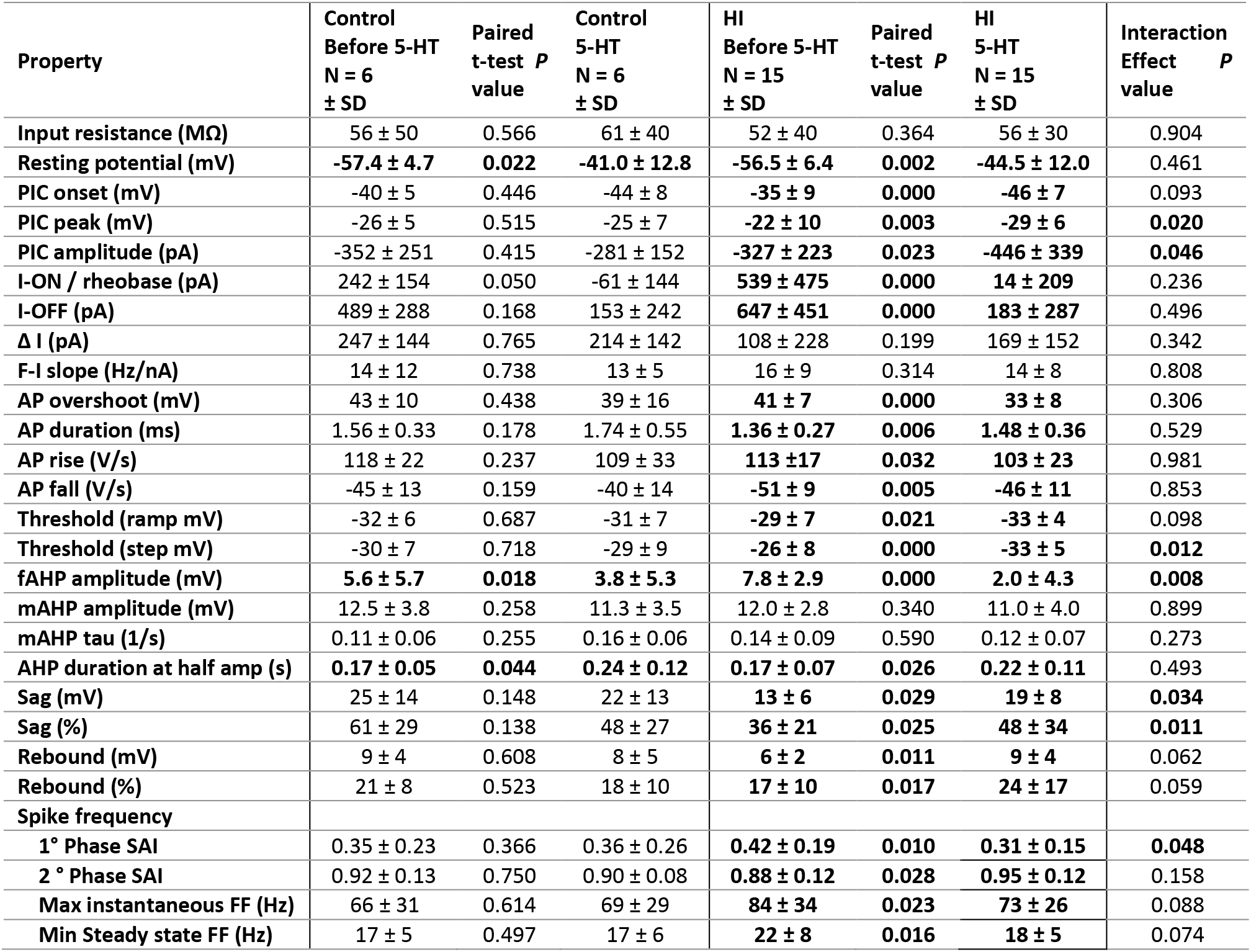
Complete results summary and statistics for control and HI MNs (all). Results from paired t-tests are included between the before and after 5-HT values for each group. The interaction P value listed in the rightmost column is from the two-way repeated measures ANOVA; P values <0.05 indicate that control and HI groups responded differently to 5-HT cocktail (interaction between surgical group and drug treatment with 5-HT cocktail). Abbreviations: PIC, persistent inward current; I-ON, current at firing onset; I-OFF, current at firing offset; ΔI, difference between I-OFF and I-ON; F-I Slope, frequency-current relationship; AP, action potential; AHP, afterspike afterhyperpolarization; fAHP, fast AHP; mAHP, medium AHP; SAI, spike adaptation index; FF, firing frequency.

| Property | Control<br>Before 5-HT<br>N = 6<br>± SD | Paired<br>t-test P<br>value | Control<br>5-HT<br>N = 6<br>± SD | HI<br>Before 5-HT<br>N = 15<br>± SD | Paired<br>t-test P<br>value | HI<br>5-HT<br>N = 15<br>± SD | Interaction<br>Effect P<br>value |
| --- | --- | --- | --- | --- | --- | --- | --- |
| Input resistance (MΩ) | 56 ± 50 | 0.566 | 61 ± 40 | 52 ± 40 | 0.364 | 56 ± 30 | 0.904 |
| Resting potential (mV) | -57.4 ± 4.7 | **0.022** | -41.0 ± 12.8 | -56.5 ± 6.4 | **0.002** | -44.5 ± 12.0 | 0.461 |
| PIC onset (mV) | -40 ± 5 | 0.446 | -44 ± 8 | -35 ± 9 | **0.000** | -46 ± 7 | 0.093 |
| PIC peak (mV) | -26 ± 5 | 0.515 | -25 ± 7 | -22 ± 10 | **0.003** | -29 ± 6 | **0.020** |
| PIC amplitude (pA) | -352 ± 251 | 0.415 | -281 ± 152 | -327 ± 223 | **0.023** | -446 ± 339 | **0.046** |
| I-ON / rheobase (pA) | 242 ± 154 | 0.050 | -61 ± 144 | **539 ± 475** | **0.000** | **14 ± 209** | 0.236 |
| I-OFF (pA) | 489 ± 288 | 0.168 | 153 ± 242 | **647 ± 451** | **0.000** | **183 ± 287** | 0.496 |
| Δ I (pA) | 247 ± 144 | 0.765 | 214 ± 142 | 108 ± 228 | 0.199 | 169 ± 152 | 0.342 |
| F-I slope (Hz/nA) | 14 ± 12 | 0.738 | 13 ± 5 | 16 ± 9 | 0.314 | 14 ± 8 | 0.808 |
| AP overshoot (mV) | 43 ± 10 | 0.438 | 39 ± 16 | **41 ± 7** | **0.000** | **33 ± 8** | 0.306 |
| AP duration (ms) | 1.56 ± 0.33 | 0.178 | 1.74 ± 0.55 | **1.36 ± 0.27** | **0.006** | **1.48 ± 0.36** | 0.529 |
| AP rise (V/s) | 118 ± 22 | 0.237 | 109 ± 33 | **113 ± 17** | **0.032** | **103 ± 23** | 0.981 |
| AP fall (V/s) | -45 ± 13 | 0.159 | -40 ± 14 | -51 ± 9 | **0.005** | -46 ± 11 | 0.853 |
| Threshold (ramp mV) | -32 ± 6 | 0.687 | -31 ± 7 | -29 ± 7 | **0.021** | -33 ± 4 | 0.098 |
| Threshold (step mV) | -30 ± 7 | 0.718 | -29 ± 9 | -26 ± 8 | **0.000** | -33 ± 5 | **0.012** |
| fAHP amplitude (mV) | **5.6 ± 5.7** | **0.018** | **3.8 ± 5.3** | **7.8 ± 2.9** | **0.000** | **2.0 ± 4.3** | **0.008** |
| mAHP amplitude (mV) | 12.5 ± 3.8 | 0.258 | 11.3 ± 3.5 | 12.0 ± 2.8 | 0.340 | 11.0 ± 4.0 | 0.899 |
| mAHP tau (1/s) | 0.11 ± 0.06 | 0.255 | 0.16 ± 0.06 | 0.14 ± 0.09 | 0.590 | 0.12 ± 0.07 | 0.273 |
| AHP duration at half amp (s) | **0.17 ± 0.05** | **0.044** | **0.24 ± 0.12** | **0.17 ± 0.07** | **0.026** | **0.22 ± 0.11** | 0.493 |
| Sag (mV) | 25 ± 14 | 0.148 | 22 ± 13 | **13 ± 6** | **0.029** | **19 ± 8** | **0.034** |
| Sag (%) | 61 ± 29 | 0.138 | 48 ± 27 | **36 ± 21** | **0.025** | **48 ± 34** | **0.011** |
| Rebound (mV) | 9 ± 4 | 0.608 | 8 ± 5 | **6 ± 2** | **0.011** | **9 ± 4** | 0.062 |
| Rebound (%) | 21 ± 8 | 0.523 | 18 ± 10 | **17 ± 10** | **0.017** | **24 ± 17** | 0.059 |
| Spike frequency |  |  |  |  |  |  |  |
| 1° Phase SAI | 0.35 ± 0.23 | 0.366 | 0.36 ± 0.26 | **0.42 ± 0.19** | **0.010** | **0.31 ± 0.15** | **0.048** |
| 2° Phase SAI | 0.92 ± 0.13 | 0.750 | 0.90 ± 0.08 | **0.88 ± 0.12** | **0.028** | **0.95 ± 0.12** | 0.158 |
| Max instantaneous FF (Hz) | 66 ± 31 | 0.614 | 69 ± 29 | **84 ± 34** | **0.023** | **73 ± 26** | 0.088 |
| Min Steady state FF (Hz) | 17 ± 5 | 0.497 | 17 ± 6 | **22 ± 8** | **0.016** | **18 ± 5** | 0.074 |

Consistent with changes to PICs, firing properties of HI MNs were also strongly impacted by the 5-HT cocktail. Two-way repeated measures ANOVA identified a significant interaction effect between surgical group (control or HI) and drug treatment (presence or absence of 5-HT cocktail) on voltage threshold (measured on step; (F[1,19]=7.80, *P*=0.012), indicating that 5-HT does not have the same effect on control and HI MN excitability. Additional affected parameters included the current at firing onset (rheobase; I-ON) and offset (I-OFF) on current ramps, which were decreased by the 5-HT cocktail (I-ON, *P*=0.000; I-OFF, *P*=0.000; **Figure 2**). Again, all HI MNs were pooled because HI-affected and HI-unaffected MNs behaved similarly: I-ON was reduced in HI affected MNs (before v. after 5-HT, *P*=0.008) and HI unaffected MNs (before v. after 5-HT, *P*=0.017); I-OFF was lower in both HI unaffected (before v. after 5-HT, *P*=0.027) and HI affected (before v. after 5-HT, *P*=0.001) MNs. None of these properties were significantly changed by 5-HT cocktail application to control MNs (all *P*≥0.05). These changes demonstrate that the excitability of HI MNs increased in the presence of 5-HT cocktail.

**Figure 2:**
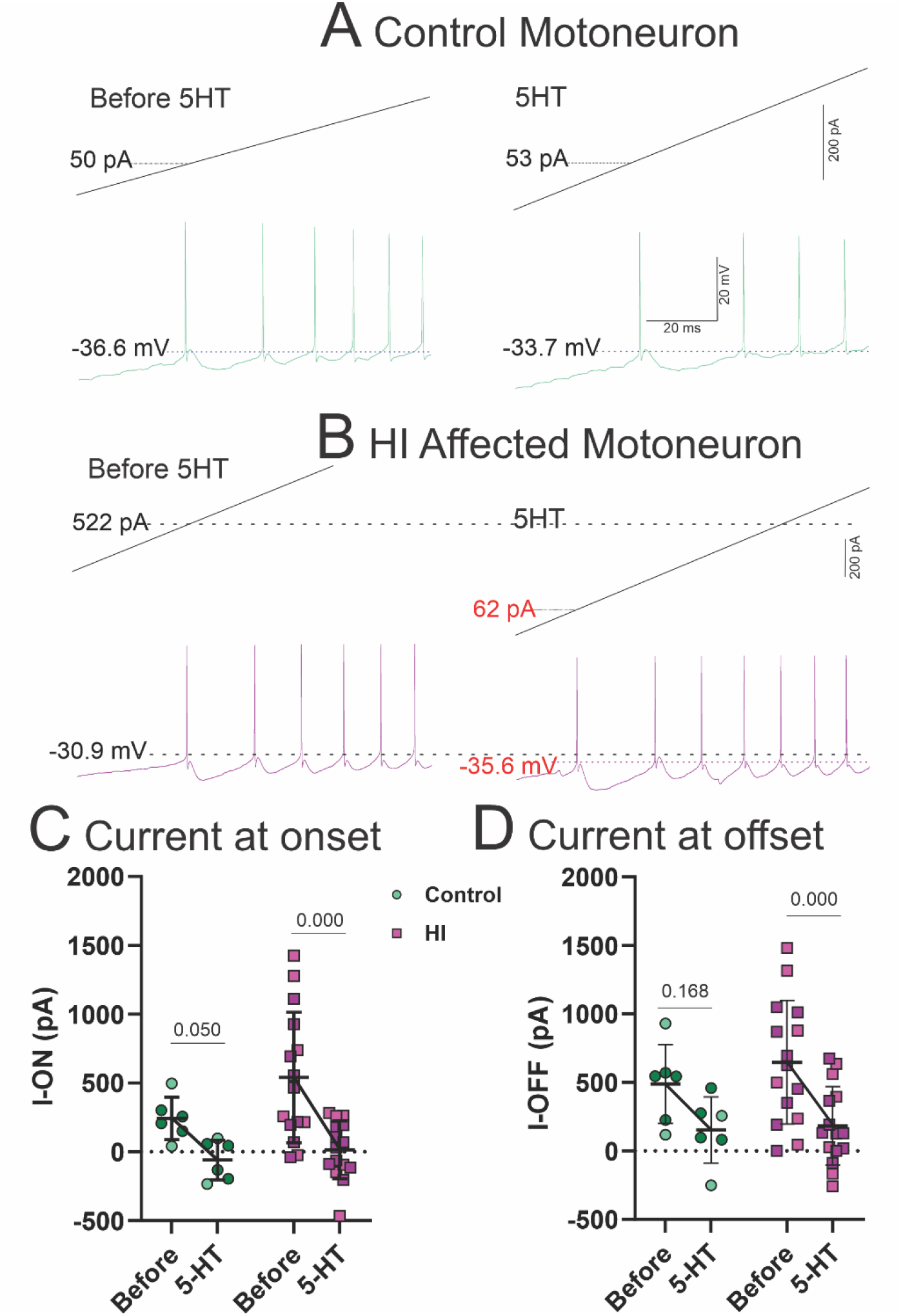
Action potential firing was differently modulated by 5-HT cocktail in control and HI MNs. (**A-B**) Examples of MN responses include a control MN showing little change with the 5-HT cocktail, while threshold voltage and current in an HI MN were lowered by the 5-HT cocktail. I-ON (rheobase) and I-OFF **(C-D)** were significantly altered by 5-HT cocktail in HI MNs but not in controls. Data are represented as mean ± SD; N=6 control MNs and N=15 HI MNs (all before and after application of 5-HT cocktail). *P* values, displayed above pairs, are based on paired t-tests (before and after application of 5-HT cocktail). Kits constituting the sham subgroup (N=4) of the control group are shaded dark green and kits constituting the motor-affected subgroup (N=9) of the HI group are shaded dark pink. Lines connecting means before and after 5-HT visually represent the interaction effect of the two-way repeated measures ANOVA. This is seen by comparing the slopes of the lines between groups.

Spike frequency adaptation (SFA) occurs when a neuron’s firing rate declines over time despite being activated by a constant suprathreshold current. As can be appreciated from the examples in **Figure 3**, the initial rate of action potential firing is typically much faster than the sustained firing rate (Shinomoto et al., 2003). Rabbit MNs often fire doublets (two rapid spikes at the onset of depolarization). These initial spikes produce the maximum instantaneous firing frequency, which is the fastest spiking a MN was observed to produce, while we measured the minimum steady state firing rate as the lowest sustained rate of firing that we observed from a MN (Steele et al., 2020). The max instantaneous firing rate was recently found to be higher in HI MNs than control MNs recorded in standard aCSF (Steele et al., 2020). Here, we found that 5-HT reduced the max instantaneous firing rate of HI MNs (before v. after 5-HT, *P*=0.023), but not control MNs (before v. after 5-HT, *P*=0.614; see Table 1). In addition, the lowest steady state firing rate observed in HI MNs was significantly reduced by 5-HT (before v. after 5-HT, *P*=0.016), but this was not the case for control MNs (before v. after 5-HT, *P*=0.497). This can be appreciated from Figure 3A-B. The control MN shows no change in firing rate upon 5-HT cocktail application, while the HI MN with a steady state firing rate of ~6 Hz before 5-HT (third step, black trace), shows a decrease in firing rate to ~4 Hz in 5-HT (first step, purple trace).

**Figure 3:**
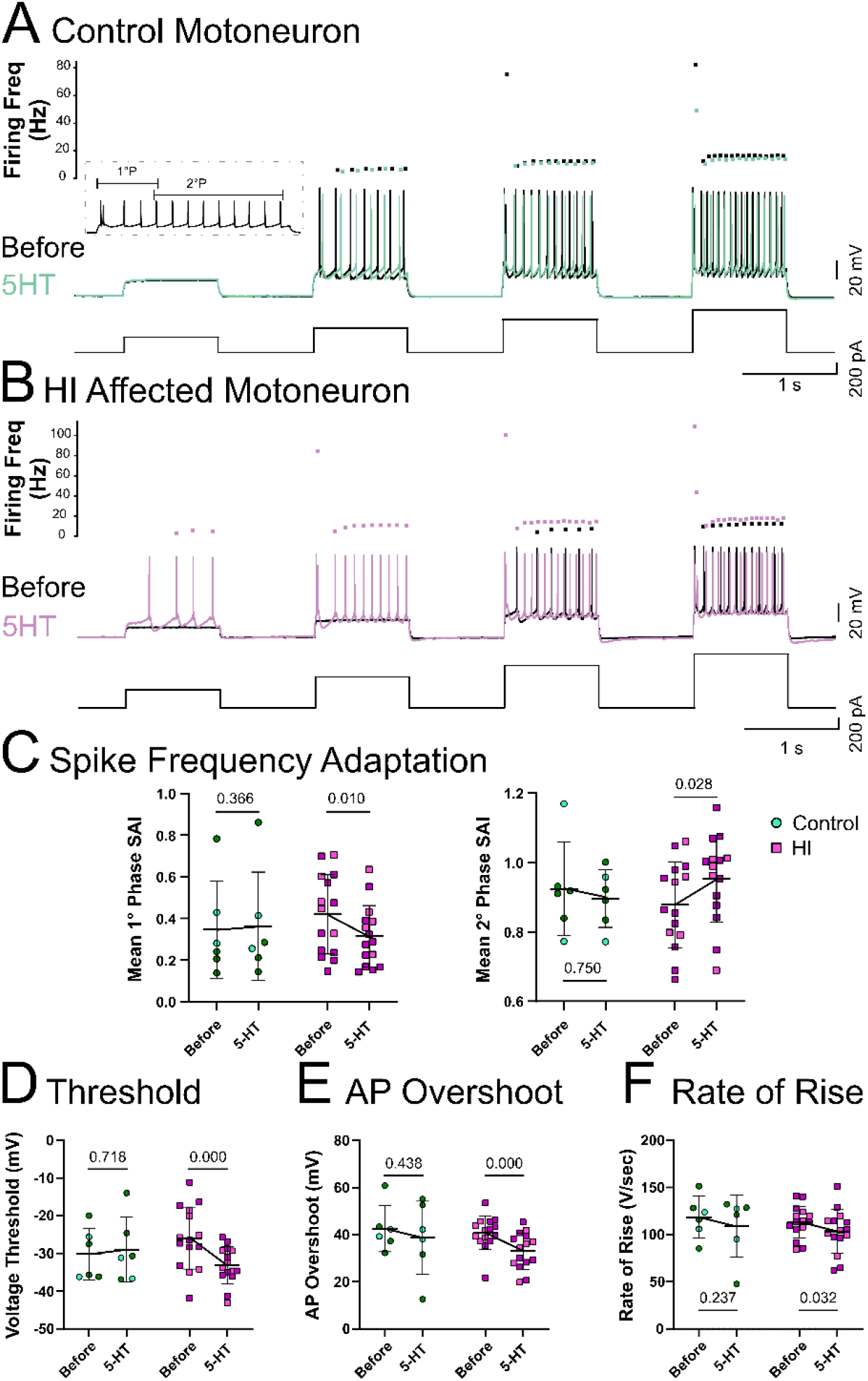
Altered firing patterns in HI MNs in the presence of 5-HT cocktail. **(A-B)** Examples of control and HI MN responses to 1 s depolarizing current steps; inset visually defines primary and secondary SAI phases. 5-HT cocktail did not change firing in control MNs (black and green squares, before and after 5-HT cocktail, respectively) but reduced minimum steady state firing frequency in HI MNs (black and pink squares before and after 5-HT cocktail, respectively). **(C)** Primary phase SAI was reduced in HI MNs after application of 5-HT cocktail, demonstrating more SFA. However, there was also an increase in the secondary phase SAI (less deceleration) in HI MNs in the presence of 5-HT cocktail. Changes in firing patterns occurred alongside changes in properties indicative of altered voltage-gated Na^+^ channel conductance: **(D)** voltage threshold, **(E)** action potential overshoot and **(F)** action potential rate of rise were all reduced in HI MNs after 5-HT cocktail. Data are represented as mean ± SD; N=6 control MNs and N=15 HI MNs (all before and after application of 5-HT cocktail). *P* values, displayed above pairs, are based on paired t-tests (before and after application of 5-HT cocktail). Kits constituting the sham subgroup (N=4) of the control group are shaded dark green and kits constituting the motor-affected subgroup (N=9) of the HI group are shaded dark pink. Lines connecting means before and after 5-HT visually represent the interaction effect of the two-way repeated measures ANOVA. This is seen by comparing the slopes of the lines between groups. Abbreviations: SAI, spike accommodation index; AP, action potential.

In order to quantify the rate of change within a single step, we also measured spike accommodation. The first 5 spikes in a step compose the primary phase (1°P SAI=ISI_First_/ISI_4th_) of firing while the secondary phase (2°P) comprises the next few hundreds of milliseconds from the 5^th^ spike to the last spike on the step (2°P SAI=ISI_5th_/ISI_Last_; Figure 3 inset). Other studies, which have used longer current pulses, have referred to these phases as the initial and early phases, respectively, which are followed by a late phase comprising another period of tens of seconds to minutes (Granit et al., 1963; Kernell, 1965; Kernell and Monster, 1982; Smith and Brownstone, 2022). Here we focused on only primary and secondary phases, using 1 s current pulses of increasing amplitude before and after application of the 5-HT cocktail. Two-way repeated measures ANOVA identified a significant interaction effect between surgical group (control or HI) and drug treatment (presence or absence of 5-HT cocktail) on mean primary phase SAI (F[1,19]=4.45, *P*=0.048), indicating prominent deceleration of the firing rate in the initial 5 spikes in HI MNs in 5-HT but a different effect (lack of response) in control MNs (Figure 3C, left panel). Interestingly, we found a significant increase in secondary phase SAI in HI MNs upon application of 5-HT (before v. after 5-HT, *P*=0.028), indicating less deceleration later in the step, including late acceleration in some HI MNs, but not control MNs (before v. after 5-HT, *P*=0.750; Figure 3B last step and 3C, right panel). In other words, while firing rate deceleration was more pronounced in the primary phase of HI MNs in the 5-HT cocktail (purple traces/dots compared to black traces/dots in Figure 3B), the secondary phase showed less deceleration in HI MNs in the 5-HT cocktail. This can be appreciated from the frequency plots above the firing examples in Figure 3A-B, where firing rate actually accelerates slightly in the secondary phase of firing in the HI MN in the 5-HT cocktail, particularly the last step, while the control MN shows no change in firing frequency with 5-HT. Because the primary and secondary phases of SFA are driven by different conductances, Ca^2+^ and Na^+^, respectively (read more in the discussion), changes in SFA occur alongside changes in other properties that are also affected by these currents, including the voltage threshold, AP height or overshoot, and the rate of rise of the action potential (Figure 3D-F), all properties that depend heavily on voltage gated Na^+^ channels. In the presence of the 5-HT cocktail, there were significant changes in all of these properties in HI, but not control MNs. Modulation of repetitive firing characteristics in MNs could contribute to weakness or central fatigue (Rose and McGill, 2005; Smith and Brownstone, 2022).

Within litters of rabbits exposed to HI, there is variability in the degree of motor dysfunction from one rabbit kit to the next. Which factors determine motor dysfunction or function are naturally of great interest. Application of the 5-HT cocktail elicited almost identical effects in MNs from HI kits that were motor affected as in MNs from kits that were HI unaffected, as mentioned previously and detailed in **Table 2**. However, HI affected MNs and HI unaffected MNs did show a different response to the 5-HT cocktail in one particular property: post-inhibitory rebound (also known as sag/I_H_) was significantly augmented by the 5-HT cocktail in HI affected MNs but not HI unaffected MNs (**Figure 4**). Specifically, the amplitude of the sag potential exhibited a 75% increase in HI affected MNs in the presence of 5-HT cocktail, while that of HI unaffected MNs was non-significant (HI affected MNs before v. after 5-HT*, P*=0.045; HI unaffected MNs before v. after 5-HT, *P*=0.436). The related post-inhibitory rebound potential also significantly increased in response to application of the 5-HT cocktail in HI affected but not HI unaffected MNs (HI affected MNs before v. after 5-HT, *P*=0.045; HI unaffected MNs before v. after 5-HT, *P*=0.183). Larger I_H_-mediated sag and post-inhibitory rebound in the HI affected MNs would increase the likelihood of action potentials occurring spontaneously after synaptic inhibition, such as during a stretch-reflex-induced spasm. This suggests that specific ionic conductances are unique to MNs in kits with motor dysfunction (more in the discussion).

**Table 2:** Comparison of HI affected and unaffected MNs. Results from paired t-tests are included between the before and after 5-HT values for each group. The interaction P value listed in the rightmost column is from the two-way repeated measures ANOVA; P values <0.05 indicate that control and HI groups respond differently to 5-HT cocktail (interaction between group and treatment effect of 5-HT cocktail). Abbreviations: PIC, persistent inward current; I-ON, current at firing onset; I-OFF, current at firing offset; ΔI, difference between I-OFF and I-ON; F-I Slope, frequency-current relationship; AP, action potential; AHP, afterspike afterhyperpolarization; SAI, spike accommodation index; FF, firing frequency.

| Property | HI unaffected before 5-HT<br>N = 6<br>± SD | Paired T test p value | HI unaffected 5-HT<br>N = 5<br>± SD | HI affected Before 5-HT<br>N = 9<br>± SD | Paired T test p value | HI affected 5-HT<br>N = 15<br>± SD | Interaction Effect P value |
| --- | --- | --- | --- | --- | --- | --- | --- |
| Input resistance (MΩ) | 40 ± 14 | 0.019 | 49 ± 16 | 60 ± 50 | 0.933 | 61 ± 37 | 0.347 |
| Resting potential (mV) | -58.0 ± 8.2 | 0.118 | -46.3 ± 13.9 | -55.5 ± 5.1 | 0.010 | -43.4 ± 11.4 | 0.954 |
| PIC onset (mV) | -36 ± 9 | 0.006 | -49 ± 9 | -35 ± 10 | 0.010 | -44 ± 5 | 0.327 |
| PIC Peak (mV) | -22 ± 12 | 0.044 | -29 ± 9 | -22 ± 10 | 0.043 | -28 ± 4 | 0.721 |
| PIC Amplitude (pA) | -322 ± 187 | 0.234 | -439 ± 215 | -330 ± 256 | 0.066 | -450 ± 415 | 0.980 |
| I-ON / rheobase (Pa) | 648 ± 600 | 0.017 | 17 ± 305 | 466 ± 393 | 0.008 | 12 ± 137 | 0.424 |
| I-OFF (pA) | 742 ± 582 | 0.027 | 199 ± 382 | 583 ± 364 | 0.001 | 172 ± 228 | 0.457 |
| Δ I (pA) | 94 ± 83 | 0.085 | 182 ± 137 | 117 ± 294 | 0.564 | 160 ± 169 | 0.647 |
| F-I Slope (Hz / nA) | 16 ± 10 | 0.534 | 15 ± 9 | 16 ± 10 | 0.396 | 13 ± 9 | 0.614 |
| AP Overshoot (mV) | 41 ± 3 | 0.015 | 33 ± 8 | 41 ± 9 | 0.006 | 33 ± 8 | 0.778 |
| AP duration (ms) | 1.31 ± 0.30 | 0.025 | 1.42 ± 0.30 | 1.39 ± 0.26 | 0.068 | 1.52 ± 0.41 | 0.853 |
| AP rise (V/s) | 112 ± 15 | 0.254 | 103 ± 15 | 114 ± 19 | 0.090 | 104 ± 28 | 0.896 |
| AP fall (V/s) | -52 ± 8 | 0.091 | -47 ± 7 | -50 ± 10 | 0.036 | -46 ± 14 | 0.595 |
| Threshold (ramp mv) | -29 ± 8 | 0.061 | -33 ± 5 | -30 ± 6 | 0.195 | -32 ± 3 | 0.424 |
| Threshold (step mv) | -26 ± 7 | 0.009 | -34 ± 5 | -26 ± 9 | 0.011 | -33 ± 5 | 0.956 |
| fAHP Amplitude (mV) | 9.2 ± 1.7 | 0.013 | 3.3 ± 3.7 | 6.9 ± 3.2 | 0.001 | 1.1 ± 4.8 | 0.930 |
| mAHP Amplitude (mV) | 10.1 ± 2.3 | 0.576 | 9.3 ± 3.8 | 13.3 ± 2.4 | 0.471 | 12.2 ± 3.8 | 0.882 |
| mAHP tau (1/s) | 0.14 ± 0.09 | 0.941 | 0.13 ± 0.10 | 0.14 ± 0.10 | 0.532 | 0.12 ± 0.05 | 0.749 |
| AHP duration at half amp (s) | 0.16 ± 0.05 | 0.226 | 0.22 ± 0.14 | 0.18 ± 0.08 | 0.032 | 0.22 ± 0.10 | 0.574 |
| Sag (mV) | 14 ± 7 | 0.436 | 16 ± 6 | 12 ± 6 | 0.045 | 21 ± 9 | 0.236 |
| Sag (%) | 32 ± 13 | 0.540 | 36 ± 15 | 38 ± 26 | 0.031 | 57 ± 40 | 0.140 |
| Rebound (mV) | 6 ± 2 | 0.183 | 9 ± 4 | 6 ± 3 | 0.045 | 10 ± 4 | 0.660 |
| Rebound (%) | 15 ± 6 | 0.215 | 20 ± 11 | 19 ± 12 | 0.055 | 27 ± 20 | 0.546 |
| Spike frequency |  |  |  |  |  |  |  |
| 1° Phase SAI | 0.48 ± 0.15 | 0.166 | 0.39 ± 0.14 | 0.38 ± 0.21 | 0.043 | 0.27 ± 0.14 | 0.828 |
| 2° Phase SAI | 0.90 ± 0.10 | 0.603 | 0.93 ± 0.12 | 0.86 ± 0.14 | 0.024 | 0.97 ± 0.13 | 0.192 |
| Max Instantaneous FF (Hz) | 73 ± 44 | 0.341 | 63 ± 29 | 91 ± 26 | 0.025 | 79 ± 22 | 0.813 |
| Min Steady State FF (Hz) | 23 ± 10 | 0.201 | 18 ± 5 | 22 ± 7 | 0.041 | 18 ± 4 | 0.773 |

**Figure 4:**
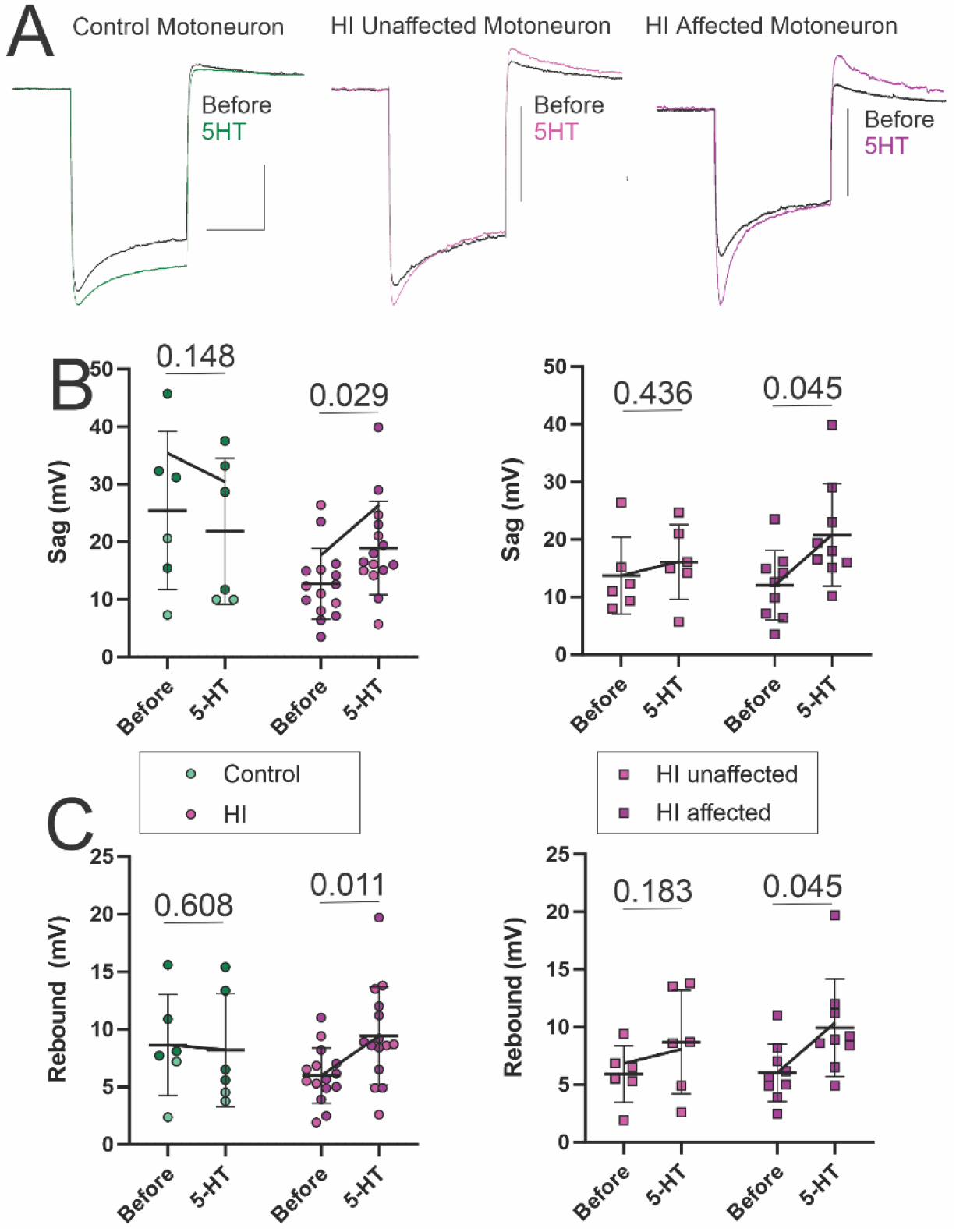
Hyperpolarization-activated sag and post-inhibitory rebound were altered most prominently in HI affected MNs (**A**). The amplitudes of sag **(B)** and post-inhibitory rebound **(C**) potentials in response to hyperpolarizing current injection were increased in HI but not control MNs (left panels). HI MNs are replotted separately (right panels) to show the different responsiveness of these properties to modulation by 5-HT in HI affected v. HI unaffected MNs. Scale bars in A: horizontal = 500 ms (scale applies to all traces); vertical = 20 mV in all traces. Kits constituting the sham subgroup (N = 4) of the control group are shaded dark green and kits constituting the motor-affected subgroup (N = 9) of the HI group are shaded dark pink. Data are represented as mean ± SD; N=6 control MNs and N=15 HI MNs (all before and after application of 5-HT cocktail). Data are represented as mean ± SD. *P* values, displayed above pairs, are based on paired t-tests (before and after application of 5-HT cocktail). Lines connecting means before and after 5-HT visually represent the interaction effect of the twoway repeated measures ANOVA. This is seen by comparing the slopes of the lines between groups.

To test the hypothesis that downregulation of the inhibitory 5-HT_1A_ receptor in HI MNs contributes to their altered sensitivity to serotonergic modulation, we performed immunohistochemistry for 5-HT_1A_ receptors in lumbar spinal MNs at P5 (**Figure 5**). We determined that significantly fewer MNs (35.3%; a 26.8% reduction) expressed the 5-HT_1A_ receptor in HI rabbit spinal cords compared to age-matched control MNs (48.2%; *P*=0.022). Interestingly, MNs from HI affected kits appeared to have the lowest relative expression of the 5-HT_1A_ receptor. Thus, reduced expression of 5-HT_1A_ receptors in HI MN populations may contribute to hyperexcitability in the presence of 5-HT.

**Figure 5:**
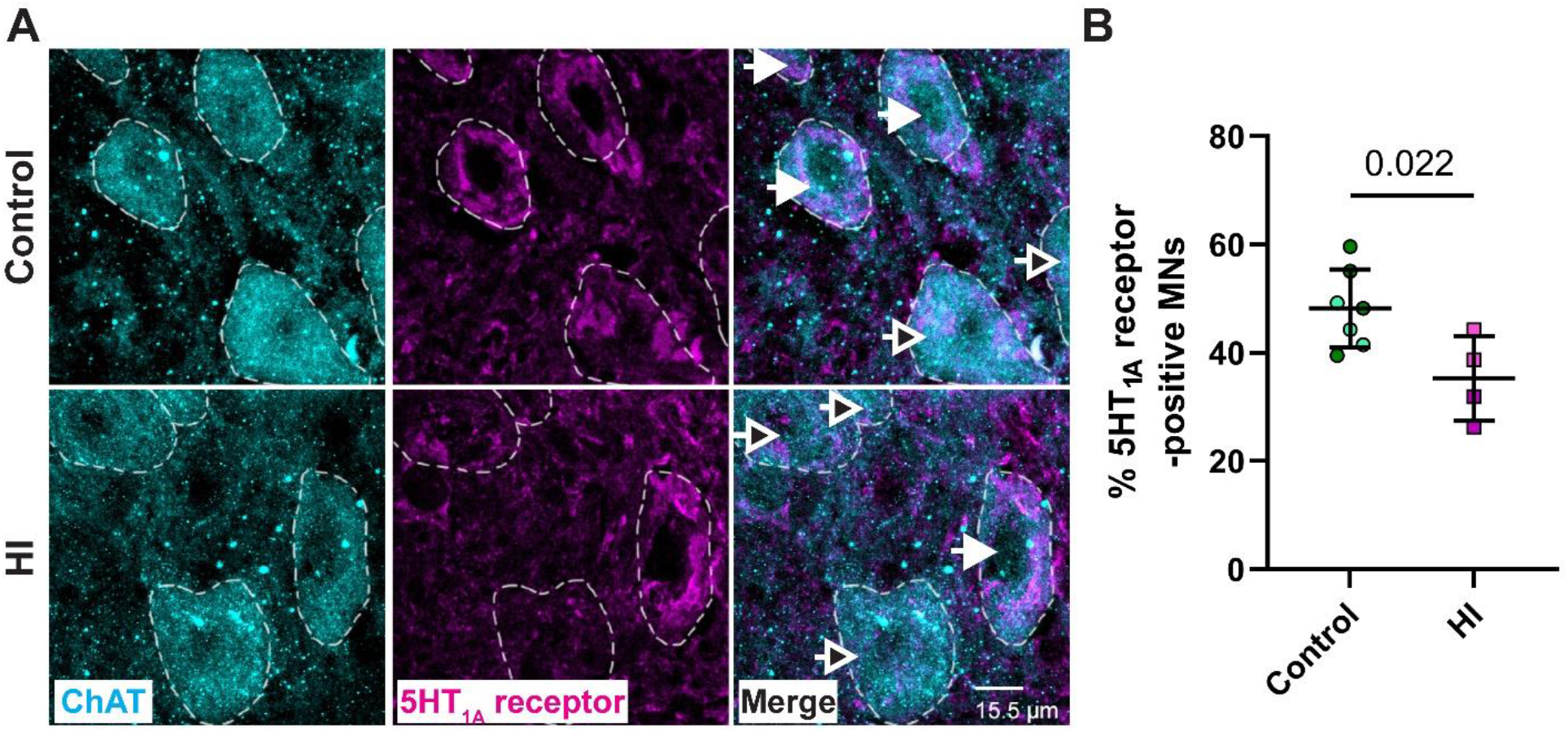
Relatively fewer MNs expressed the 5-HT_1A_ receptor in HI rabbit spinal cords at P5 compared to age-matched controls. A: Representative maximum intensity z-projections of lower lumbar MNs from control (top) and HI (bottom) rabbits immunostained for the MN marker choline acetyltransferase (ChAT; pseudocolored cyan, left) and the 5-HT_1A_ receptor (pseudocolored magenta, middle; merged images on right). MN somas are traced with dashed white lines; white arrows indicate 5-HT_1A_ receptor-positive MNs; black-filled arrows indicate 5-HT_1A_ receptor-negative MNs. B: Quantification of the relative abundance (percent) of 5-HT_1A_ receptor-positive MNs for control and HI kits. Kits constituting the sham subgroup of the control group are shaded dark green and kits constituting the motor-affected subgroup of the HI group are shaded dark pink. A significantly smaller percentage of lower lumbar MNs expressed the 5-HT_1A_ receptor in the spinal cord of HI rabbits (mean: 35.3%) compared to controls (mean: 48.2%) at P5 (\**P*=0.022, unpaired t-test). Data are represented as mean ± SD; N=4-7 rabbits per group; n=131-729 total MNs per rabbit.

## Discussion

### Summary

In this study, we demonstrate that the electrophysiological response of neonatal rabbit spinal MNs to 5-HT is altered by prenatal HI. Specifically, MNs exposed to prenatal HI were primed to become hyperexcitable in response to the 5-HT cocktail, as shown through hyperpolarization of the PIC and threshold voltage for action potentials. HI MNs exposed to the 5-HT cocktail exhibited a reduction in maximum firing frequencies yet had more sustained firing in the secondary phase of firing while control MNs did not show significant changes in PICs, threshold voltage, or firing patterns in the presence of 5-HT. We determine that fewer HI MNs expressed the 5-HT_1A_ receptor, an inhibitory G-protein coupled receptor that is important in central fatigue, compared to control MNs. Downregulation of 5-HT_1A_ receptors in HI MNs together with altered firing patterns (increased secondary phase SAI) in 5-HT show that HI MNs are more fatigue-resistant. These properties likely contribute to hypertonia observed in motor affected HI rabbit kits, and more broadly, in spastic CP.

### Implications for motor dysfunction

The main finding of this study was that HI MNs responded to exogenously applied 5-HT with a robust increase in excitability, due at least in part to their decreased expression of 5-HT_1A_ receptors compared to control MNs. These inhibitory G protein-coupled receptors are coupled to inhibitory G_i_/G_o_ proteins (Barnes and Sharp, 1999), and mediate reduced excitability and central fatigue (Cotel et al., 2013; D’Amico et al., 2017). Loss of 5-HT_1A_ receptors and changes in SFA (particularly the reduction in secondary phase SAI) could both be directly related to the increased resistance to fatigue that is observed in muscles of people with CP (Moreau et al., 2008, 2009). Increased fatigue resistance correlates with lower mobility in patients (Moreau et al., 2009), suggesting this is a serious problem leading to motor dysfunction.

Neonatal rabbit kits that experienced prenatal HI injury vary in degree of motor dysfunction. Here, we found that MNs from motor-affected and motor-unaffected HI kits had similar responses to modulation by 5-HT, demonstrating that the response to 5-HT alone is not sufficient to explain motor dysfunction after prenatal HI. Rather, *it is the presence of elevated spinal 5-HT* in motor-affected HI kits, which was demonstrated previously (Drobyshevsky et al., 2015), that separates those kits that are affected with motor dysfunction from those that are spared (HI unaffected). Electrophysiological abnormalities specific to MNs of affected HI kits include a more pronounced increase in I_H_ currents in the presence of 5-HT, which manifests as larger sag and rebound potentials. This could contribute to initiation of MN activity following inhibition, and as described previously, prolonged firing with diminishing current injection (reduced ΔI), depolarized resting membrane potential and longer dendritic length (Steele et al., 2020). An important takeaway from this study is that both altered spinal 5-HT and altered MN excitability contribute to motor dysfunction in CP.

### Contribution of 5-HT to MN excitability

HI rabbit MNs responded to the 5-HT cocktail with hyperpolarization of the PIC and voltage threshold, increased I_H_, and altered patterns of firing. These changes are generally consistent with previous studies in neonatal mice and rats (Wang and Dun, 1990; Elliot and Wallis, 1992; Ziskind-Conhaim et al., 1993). The response of control MNs to 5-HT appears modest compared to HI MNs, however, consistent with previous studies in neonatal MNs, control rabbit MNs depolarized by an average of 16 mV in the presence of 5-HT (resting potential before v. during 5-HT, *P*=0.022; Table 1) (Wang and Dun, 1990; Ziskind-Conhaim et al., 1993). Serotonergic enhancement of PICs, observed here in HI but not control MNs, has been previously demonstrated in adult but not neonatal MNs (Larkman and Kelly, 1992; Bayliss et al., 1995; Li et al., 2006). Neuromodulation of the PIC by 5-HT (and other neuromodulators) becomes possible with increasing postnatal age (Revill et al., 2019). Increased PICs also impact firing properties. Decreased adaptation of the firing rate in HI MNs in the secondary phase (reflecting not quite acceleration, but less deceleration of the firing rate of HI MNs in response to 5-HT cocktail) could be due to the increased amplitude of the PIC observed in HI but not control MNs (Granit et al., 1963; Kernell, 1965; Kernell and Monster, 1982; Smith and Brownstone, 2022). Consistent with this result, serotonergic neuromodulation of spinal MNs has been found previously to reduce SFA (Martin, 2002; McClelland and Parker, 2017), essentially prolonging sustained firing. Overall, serotonin contributes to a robust increase in excitability in HI MNs.

### Effects of 5-HT on ionic conductances in HI MNs

Increased MN excitability in the presence of 5-HT is mediated by neuromodulation of ionic conductances, including voltage-gated Na^+^ and Ca^2+^ channels (Heckman et al., 2009). SFA, PICs, and threshold all depend on voltage-gated Na^+^ and Ca^2+^ channels. Previous studies have found activation of 5-HT_2A/2B/2C_ receptors upregulate both Na^+^ and Ca^2+^ PICs and increase intrinsic excitability (Ladewig et al., 2004; Harvey et al., 2006a, 2006b). Effects of 5-HT on SFA in HI MNs were mixed, likely due to differential effects of 5-HT on Na^+^ and Ca^2+^ channels. The primary phase of firing was dominated by the presence of the initial doublet that is Ca^2+^-dependent (Kobayashi et al., 1997; Powers et al., 1999). Since inhibition of high voltage-gated Ca^2+^ channels by 5-HT has been previously shown to depend on 5-HT_1A_ receptor activation (Bayliss et al 1995), it is puzzling that there is more adaptation in the firing rate in the primary phase of HI MNs despite less expression of 5-HT_1A_ receptors. The increased SAI in the secondary phase may reflect reduced inactivation of Na^+^ channels in the presence of 5-HT (Miles et al., 2005), which is also supported by the hyperpolarization in the threshold voltage in HI MNs. Increased I_H_ underlying the larger sag and rebound in HI affected MNs in 5-HT is likely due to a depolarized voltage dependence of I_H_ (Kjaerulff and Kiehn, 2001), enhancing nonspecific cation currents via HCN channels, possibly along with increased conductance in Ca_v_3 Ca^2+^ channels (Zemankovics et al., 2010; Canto-Bustos et al., 2014). In both control and HI MNs, 5-HT also reduced the fast AHP (fAHP) amplitude (control MNs before v. after 5-HT, *P*=0.018; HI MNs before v. after 5-HT, *P*=0.000; Table 1). In some cases, the fAHP was dissipated entirely after application of serotonergic drugs. This could be due to inhibition of K_IR_ current by 5-HT (Kjaerulff and Kiehn, 2001), or diminished driving force of voltage gated K^+^ currents, since 5-HT depolarized control and HI MNs and hyperpolarized the spike threshold in HI MNs. Decreased activity of K^+^ channels that repolarize the spike in HI MNs and produce the fAHP could also be the cause of the decrease in maximum instantaneous firing rates, since they permit deinactivation of Na^+^ channels needed for subsequent spikes (Jaffe and Brenner, 2018). In agreement with previous studies, many different conductances in MNs are sensitive to modulation by 5-HT, and likely depend on the specific complement of 5-HT receptors that are present.

### 5-HT receptor subtypes

We chose to use α-methyl-5-HT to study MN responses to serotonergic neuromodulation since it is a broad agonist of 5-HT_1_ and 5-HT_2_ receptors, binding to 5-HT_1B_, 5-HT_1D_, 5-HT_1F_, 5-HT_2A_, 5-HT_2B_, 5-HT_2C_ AND 5-HT_4_ receptors. More specifically, at the concentration used in this study, it is likely to mediate its effects through binding to 5-HT_2C_ and possibly 5-HT_2B_ receptors (Ismaiel et al., 1990; Lucas-Osma et al., 2019). In addition to α-methyl-5-HT, we also included citalopram in our drug cocktail because it prevents reuptake of endogenously released 5-HT. Endogenous 5-HT is released from serotonergic fibers in isolated spinal cords during periods of activity (Maclean et al., 1998; Pearlstein et al., 2005), as would occur during application of α-methyl-5-HT in the slices recorded here (many neurons in the spinal cord are brought to threshold by α-methyl-5-HT). Endogenously released 5-HT could bind to any 5-HT receptors present in the spinal cord. Thus, the experiments here show responsiveness of neonatal rabbit MNs to α-methyl-5-HT via 5-HT1 and 5-HT_2_ receptors and very possibly other 5-HT receptor subtypes to endogenous 5-HT. MNs typically express receptors from each of the seven families of 5-HT receptors (5-HT1 – 5-HT7), with the apparent exception only of 5-HT_6_ receptors (Ziskind-Conhaim et al., 1993; Morales et al., 1999; Xu et al., 2007; Cotel et al., 2013; Suwa et al., 2014; Cabaj et al., 2017). We show here that approximately half of typically developing lumbar spinal MNs express 5-HT_1A_ receptors at P5, but significantly fewer lumbar spinal MNs express this receptor after prenatal HI injury (35.3% of HI MNs v. 48.2% control MNs). Expression of 5-HT_1A_ receptors in rabbits at this age may be more widespread than in rodents. Based on electrophysiological evidence, 9% of neonatal rat MNs possess enough 5-HT_1A_ receptors to mediate a hyperpolarizing response to 5-HT. 5-HT_1A_ receptors are also expressed by human spinal MNs (Perrin et al., 2011; D’Amico et al., 2017). During postnatal development, expression of 5-HT_1A_ receptors in MNs decreases (Talley et al., 1997; Talley and Bayliss, 2000) while expression of 5-HT_2A_ and 5-HT_2C_ receptors increases (Volgin et al., 2003). Since both control and HI MNs in this study were examined at P5, age would not contribute to the results *perse*. However, HI could impact developmental processes that regulate the changing expression of 5-HT receptors. Interestingly, other factors can also alter expression of 5-HT receptors: after exercise, 5-HT_1A_ receptor mRNA decreased in MNs (Woodrow et al., 2013), while 5-HT immunoreactivity increased (Behan et al., 2012). Since activation of 5-HT_1A_ receptors causes membrane hyperpolarization and central fatigue, reduced expression of this receptor type in HI MN pools may contribute to hyperexcitability and fatigue resistance (Cotel et al., 2013). But lack of central fatigue is likely only part of the story. Evidence for increased signaling through excitatory 5-HT receptors in CP includes case reports of increased spasticity in individuals with CP after starting selective serotonin reuptake inhibitor treatment for depression (Rone and Ferrando, 1996), and reports of successful treatment of baclofen withdrawal symptoms with serotonergic antagonists (Meythaler et al., 2003; Saveika and Shelton, 2007), indicating increased expression or activation of 5-HT_2_ receptors, along with diminished 5-HT_1_ receptors, may both contribute to pathology. Our findings support further exploration of interventions for CP that target serotonergic modulation of spinal MNs and therapies to restore inhibitory/excitatory balance within spinal circuits for alleviation of spasticity.

### A role for 5-HT in etiology of motor dysfunction

Several conditions including stroke, spinal cord injury and traumatic brain injury alter serotonergic signaling, and perhaps in contrast to CP, augmenting 5-HT could improve recovery for these conditions (Murray et al., 2010, 2011; D’Amico et al., 2013; Liepert, 2016; Gu and Wang, 2018). The results of the current study suggest that diminishing spinal 5-HT or blocking 5-HT receptors on spinal MNs could be therapeutic in CP. In contrast, acute intermittent hypoxia, which acts through activation of 5-HT receptors in phrenic MNs (Tadjalli and Mitchell, 2019; Wen et al., 2019), has been found to be therapeutic for some of the above conditions. After spinal cord injury, neurons become hypersensitive to 5-HT (Husch et al., 2012), which is reminiscent of the increased responsivity of HI MNs to 5-HT that we observed in this study. Future work will focus on determining the receptor identity that mediates this robust response in HI rabbit MNs. Perhaps whether 5-HT is therapeutic or detrimental is determined by how 5-HT impacts MN excitability, and when. If 5-HT increases MN firing during a developmentally sensitive period (as it does in prenatal HI rabbits), it could permanently alter neuromuscular development, specifically the development of motor unit types. Reduced rate modulation of motor units, as indicated by changes in SAI, has been observed in CP patients (Rose and McGill, 2005). Sustained changes to MN firing frequencies could drive alterations in muscle physiology and lead to weakness and spasticity. It has been traditionally believed that hypertonia results from overactive MNs due to disinhibition (Sanger, 2003; Deon and Gaebler-Spira, 2010; Volpe et al., 2017), however it could also be due to tonic recruitment of lower threshold motor units (containing type 1 fibers), which over time leads to disuse atrophy of type 2 fibers and simultaneously stiff but weak muscles. Thus, there could be specific effects of both hypoxia and injury to the central nervous system that promote upregulation of serotonergic fiber sprouting and 5-HT receptor expression.

### Conclusion

Exploring increased serotonergic signaling in the spinal cord coincident with loss of descending corticospinal connectivity should be pursued in future studies. It is likely that preventing pathological augmentations of spinal 5-HT would be sufficient to reduce motor dysfunction after prenatal HI injury. Determining which 5-HT receptor subtypes mediate spinal MN hyperexcitability could ultimately lead to serotonin-based treatments that improve outcomes for individuals with CP.

## Data Availability Statement

All raw data (recordings and images) from which graphical and/or tabular summary data is generated is archived and fully available to The Journal upon reasonable request.

## Author Contributions

EJR, LTG, PRS, EMA, and LD contributed to acquisition, analysis and interpretation of data and drafting/revising it critically for important intellectual content. AD contributed to conception and design of the work and drafting/revising it critically for important intellectual content. MM contributed to analysis and interpretation of data and drafting/revising it critically for important intellectual content. KAQ contributed to conception and design of the work, acquisition, analysis and interpretation of the data, and drafting/revising the work for important intellectual content.

All authors approved the final version of the manuscript and agree to be accountable for all aspects of the work in ensuring that questions related to the accuracy or integrity of any part of the work are appropriately investigated and resolved; and all persons designated as authors qualify for authorship, and all those who qualify for authorship are listed.

## Acknowledgements

We thank Alyssa M. Garrett and Brianna Taylor for their help preparing z-stack photomicrographs for analysis. This research was conducted, in part, using equipment and services available through the Institutional Development Award (IDeA) Network for Biomedical Research Excellence (INBRE) under grant number P20GM103430 from the National Institute of General Medical Sciences of the National Institutes of Health.

